# Addiction-Like Severity Predicts Prolonged Oxycodone Withdrawal-Induced Allodynia in Genetically Diverse Rats

**DOI:** 10.64898/2026.05.14.725258

**Authors:** Sonja L Plasil, Lani Tieu, Chengjia Qian, Natalie Taylor, Elizabeth Sneddon, Lieselot LG Carrette, Molly Brennan, Alex Morgan, Dyar Othman, Kathleen Bai, Sara Foroutani, Giordano de Guglielmo, Marsida Kallupi, Olivier George

## Abstract

Opioid withdrawal is associated with heightened pain sensitivity, including allodynia. Although opioid-induced allodynia is well-documented in humans and animal models, the relationship between the severity of opioid withdrawal-induced allodynia and individual addiction-like behaviors remains poorly understood. To address this gap, Heterogeneous Stock rats underwent long access (12 hours/day) intravenous oxycodone self-administration, followed by measurement of mechanical sensitivity at six timepoints across three weeks of abstinence. Rats were stratified by an Addiction Index derived from individual differences in the escalation of oxycodone intake, motivation to consume oxycodone, tolerance to oxycodone’s analgesic effects, and acute withdrawal-induced mechanical pain sensitivity. Here, we show that oxycodone withdrawal induces significant and prolonged allodynia for up to three weeks, with High Addiction Index rats exhibiting greater intensity and longer duration of pain sensitivity than Low Addiction Index rats. Results remained consistent even when excluding allodynia from the Addiction Index, highlighting the robustness of the association between addiction-like severity and protracted allodynia. Linear regression associations revealed that self-administration behaviors, particularly oxycodone intake escalation and motivation to seek oxycodone, predicted subsequent withdrawal-induced allodynia severity. These findings demonstrate that greater addiction-like severity is associated with more intense and prolonged withdrawal-induced pain, supporting mechanical allodynia as a marker of addiction severity. These results motivate future work to define the mechanisms linking addiction severity to protracted opioid withdrawal-induced pain, with the goal of informing targeted clinical interventions for individuals most susceptible to severe abstinence-related allodynia.

## Introduction

The opioid epidemic poses a major public health crisis^1–4^, driven in part by the widespread misuse of prescription opioids^5–17^ such as oxycodone. Chronic opioid misuse leads not only to compulsive drug seeking but also to significant alterations in pain sensitivity^18–22^, including opioid withdrawal-induced hyperalgesia^21,23–34^ and mechanical allodynia^26,35–42^. These are major drivers of negative states associated with opioid use^5,17,43,44^. Pain states that emerge during opioid withdrawal are believed to increase relapse vulnerability^39,43–45^, yet their behavioral correlates remain incompletely understood. Further, the relationship between pain and opioid self-administration is inconsistent across preclinical studies. While chronic inflammatory pain can increase opioid intake via analgesic negative reinforcement^46^, experimental pain manipulations fail to enhance opioid seeking^47^ or reduce opioid reward sensitivity and diminish opioid self-administration^48–50^. Collectively, these discrepancies indicate that opioid reinforcement is not uniformly pain-driven and may depend on additional factors, including differences in addiction-like behavioral severity.

Opioid withdrawal-induced allodynia has been observed in both humans and animal models^19,40–42,51–54^, but individual variability in the intensity and duration of this phenomenon has not been systematically linked to addiction or addition(-like) behaviors. Clinical evidence suggests opioid-induced allodynia^40^ may last from several days to several weeks during protracted abstinence^26,39,55^, particularly after long-term or high-dose opioid use^56^. Previous preclinical studies have demonstrated that opioid-intake escalation, increased motivation to self-administer opioids, and the development of tolerance are key hallmarks of addiction-like behaviors in rodents^19^. However, the protracted timeline remains unknown as well as whether the severity and duration of opioid withdrawal-induced allodynia^42^ correlate with addiction-like behavioral profiles. Specifically, the relationship between early self-administration behaviors and subsequent pain sensitivity during protracted abstinence has not been resolved.

To address this gap, the current study tested the hypothesis that addiction-like behavioral severity predicts the magnitude and duration of mechanical allodynia during oxycodone withdrawal. Genetically diverse Heterogeneous Stock rats^19,57–65^ were trained to self-administer intravenous oxycodone (intravenous self-administration; IVSA) under short- (2 hours/day) and long-access (6 hours/day), and their addiction-like behaviors were quantified based on the escalation of oxycodone intake, motivation to consume oxycodone, tolerance to oxycodone’s analgesic effects, and acute withdrawal-induced allodynia^19^. Rats were stratified using a composite Addiction Index into High and Low subpopulations of addiction-like behavior to examine individual differences in withdrawal-induced mechanical sensitivity over a three-week abstinence period that followed oxycodone self-administration. Here, we show that a history of oxycodone self-administration leads to prolonged allodynia throughout abstinence, and this relationship strengthens with the severity of the individual animal’s Addiction Index. This approach allowed for the identification of behavioral predictors of pain vulnerability in opioid withdrawal and provides insight into potential markers of addiction severity.

## Methods

### Animals

Heterogeneous Stock rats (n=19/sex; Rat Genome Database NMcwiWFsm #13673907, RRID:RGD 13673907) were obtained from Wake Forest and the University of California San Diego School of Medicine (Drs. Leah Solberg Woods and Abraham Palmer, respectively). The oxycodone self-administering cohort included n=11 rats/sex, while the age-matched drug-free control cohort included n=8 rats/sex. This outbred population was generated from eight inbred rat strains (ACI/N, BN/SsN, BUF/N, F344/N, M520/N, MR/N, WKY/N, and WN/N) and maintained for over 100 generations to maximize genetic diversity^66^. Rats from Wake Forest were shipped at 3 – 4 weeks of age and quarantined for 2 weeks, while rats from the University of California San Diego were transferred at 6 – 7 weeks of age then housed in pairs under a 12-hour light/dark cycle (lights on at 4:00 AM) in temperature (22 ± 1°C) and humidity (45 – 55%) controlled conditions. Food (PJ Noyes Company, Lancaster, NH, USA) and water were available *ad libitum*. All procedures complied with the National Institute of Health Guide for the Care and Use of Laboratory Animals and were approved by the Institutional Animal Care and Use Committee of the University of California, San Diego.

### Intravenous Catheterization

Rats were anesthetized with vaporized isoflurane (1 – 5%; Piramal Critical Care Inc., 66794-017-25) and aseptic surgery was performed to implant chronic intravenous jugular vein catheters. Flunixin (2.5 mg/kg, s.c.; Covetrus, 11695-4025-1) and Cefazolin (330 mg/kg, i.m.; Hikma, 0143-9924-90) were administered as an analgesic and antibiotic, respectively. Following puncture of the right jugular vein with a 22G needle, catheters constructed of Micro-Renathane tubing (18 cm length, 0.023-inch inner diameter, 0.037-inch outer diameter; Braintree Scientific, Braintree, MA, USA) were inserted and secured with sutures. Catheters were anchored subcutaneously with mesh and embedded in dental acrylic (Co-Oral-Ite Dental Mfg. Co., Diamond Springs, CA, USA), terminating in a 90° angle-bend guide cannula (Shenzhen Huayon Biotech Co., Ltd, Futian Dist., Shenzhen, China), that exited dorsally. The exterior of the cannulas were protected using a plastic seal and metal cap to ensure sterility of the catheter base^62,67^. Post-operative recovery lasted 7 days, during which catheters were flushed daily with heparinized saline (10 U/ml of heparin sodium; American Pharmaceutical Partners, Schaumberg, IL, USA) in 0.9% bacteriostatic sodium chloride (Hospira, Lake Forest, IL, USA) containing Cefazolin (52.4 mg/0.2 mL).

### Drugs

Oxycodone hydrochloride (National Institute on Drug Abuse, Bethesda, MD, USA) was dissolved in 0.9% sterile saline and administered intravenously (150 μg/0.1mL/kg/infusion). Doses were calculated using the molecular weight of the oxycodone hydrochloride free base salt. This dose was based on previous studies^51,61,68^ and produces significant plasma oxycodone concentrations (∼40 ng/mL)^69^.

### Intravenous Self-Administration (IVSA)

Self-administration was conducted in operant conditioning chambers (29cm × 24cm × 19.5cm; Med Associates, St. Albans, VT, USA) enclosed in lit, sound-attenuating, ventilated environmental cubicles. The front door and back wall of the chambers were constructed of transparent plastic, and the side walls were opaque metal. Each chamber was equipped with two retractable levers (active and inactive), a cue light above the active lever, and an infusion pump (PHM-100VS-2, Med Associates Inc.) connected to the catheter via flexible tubing. Sessions were initiated by lever extension and at the onset of the dark cycle. Responses on the active lever delivered an infusion of oxycodone over 6 seconds, followed by a 20-second timeout period signaled by the cue light. Inactive lever presses were recorded but had no programmed consequence. Behavioral data and pump operation were controlled using MED-PC computer software (Med Associates Inc.).

Rats underwent four short-access sessions (2 hours/session) followed by 14 long-access sessions (12 hours/session, 5 session/week) to measure escalation of oxycodone intake. Food was available *ad libitum* during sessions, and a wooden block (3 × 3 × 3 cm) was provided for environmental enrichment.

### Progressive Ratio

Rats were tested on a Progressive Ratio (PR) schedule of reinforcement at the end of each phase (once after short access and once after long access; **Figure 1**). In the PR paradigm, the number of lever presses required for each oxycodone infusion (150 μg/kg/infusion) increased progressively within each session as follows: 1, 1, 2, 2, 3, 3, 4, 4, 6, 6, 8, 8, 10, 11, 12, 13, 14, 15, 16, 17, etc. +1 until 50, 60, 70, 80, 90, 100. The breakpoint was defined as the last completed ratio before a 60-minute period without completing the next ratio, terminating the session.

**Figure 1.**
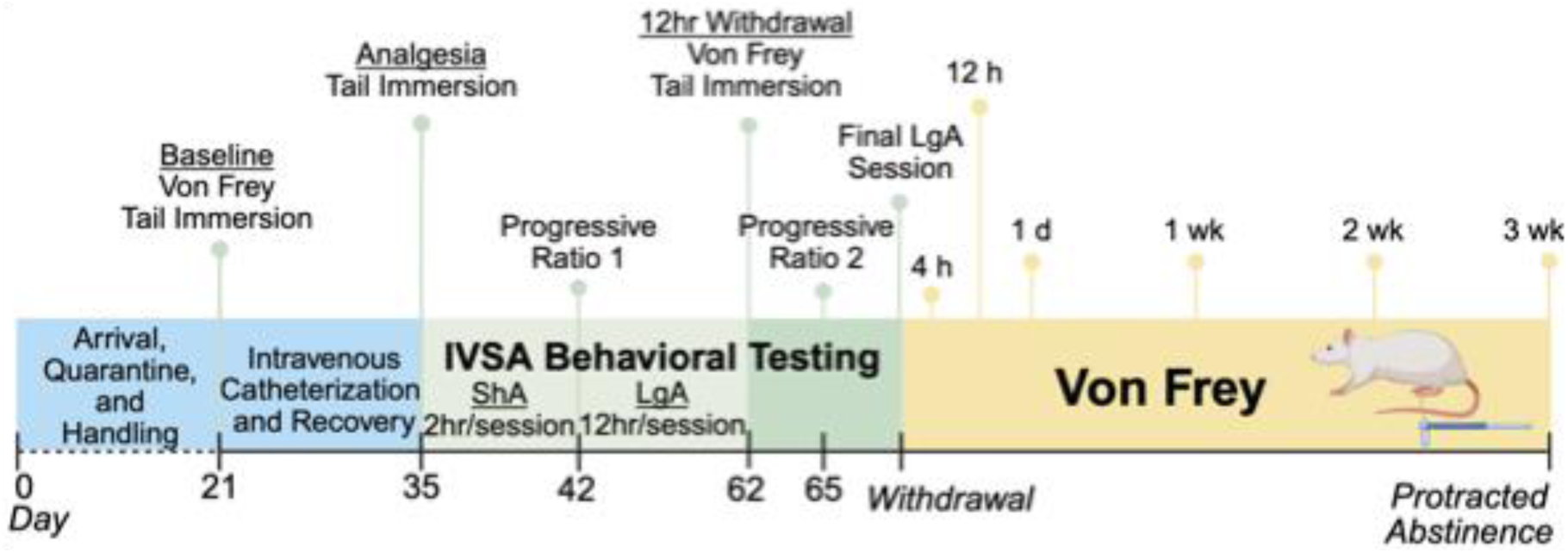
Behavioral paradigm schematic Timeline. IVSA, intravenous self-administration; ShA, Short Access; LgA, Long Access.

### Tail Immersion

Analgesia and tolerance to oxycodone’s analgesic effects were assessed in rats by measuring tail withdrawal latency from a hot water bath (52° C; Isotemp GPD 02, Fisher Scientific)^70^. Rats were gently restrained in a soft cloth pouch, and the distal 1 cm of the tail was immersed in the water bath. Latency to tail withdrawal was recorded, with a 10 second cut-off time imposed to prevent tissue damage. Measurements were obtained at three different timepoints: (1) oxycodone-naïve baseline; (2) following acute oxycodone administration (no prior drug history; 450 μg/kg i.v.); and (3) following acute oxycodone administration 12 hours into withdrawal post-long access IVSA (450 μg/kg i.v.) (**Figure 1**). A 30 second delay was imposed between i.v. infusion and tail immersion testing.

### Von Frey

Mechanical sensitivity was assessed using dynamic plantar aesthesiometer (Ugo Basile, Gemonio, Italy), adapted from previous reports^71–73^. The force required to elicit paw withdrawal was recorded three times per hind paw. Measurements were averaged across all trials for each rat. Outliers were excluded if they fell outside of 2 standard deviations from the mean. Data were expressed as the mechanical threshold in grams of force and as the percentage change from baseline. Measurements were obtained at two timepoints during the oxycodone self-administration paradigm: drug-naïve baseline and 12-hour into oxycodone withdrawal following long access IVSA. During protracted abstinence, measurements were taken at six timepoints: 4 hours, 12 hours, 24 hours, 1 week, 2 weeks, and 3 weeks-post withdrawal.

### Addiction Index

Addiction-like behavioral measures were normalized into indices based on individual Z-score’s (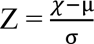), where χ is the raw value, μ is the cohort mean (per sex), and σ is the cohort standard deviation (per sex). Four behavioral metrics were normalized using sex-specific Z-scores: escalation of oxycodone intake (average intake over the last three long access sessions), motivation (LgA PR breakpoint), analgesic tolerance (tail immersion response T2-T3), and withdrawal-induced allodynia (Von Frey response). The average Z-score across all four metrics constituted the Addiction Index. Rats with a positive Addiction Index score were categorized as High Addiction Index (having high addiction-like behaviors); those with negative scores were classified as Low Addiction Index (having low addiction-like behaviors). The oxycodone self-administering cohort was therefore subdivided into n = 9 High Addiction Index (n = 4F/5M) and n = 13 Low Addiction Index (n = 7F/6M) rats. To ensure the predictive value of the Addiction Index was not simply due to the pain measure being a part of the index, we also recalculated the Addiction Index excluding the pain component. Even without the Pain Z-score, the results remained consistent, with High Addiction Index animals showing significant allodynia at 12 hours, 1 day, and 2 weeks into withdrawal (**Supplemental Figure 1A**).

### Statistical Analysis

Data were analyzed using Prism 10.0 (GraphPad, San Diego, CA). Repeated-measures Two-Way analysis of variance (ANOVA) with Bonferroni *post hoc* tests were used for longitudinal data. Group comparisons were conducted using one-tailed unpaired t-test or one-tailed Mann Whitney test for non-parametric data. Allodynia intensity was defined as the largest reduction in mechanical pain threshold observed. Allodynia duration was set as the timepoint at which allodynia had ceased; normal variability in mechanical pain threshold was estimated from Naïve animals across all timepoints, and a ±10% range was selected as the threshold for normal variability. One-Way ANOVA or Kruskal-Wallis test for non-parametric data was used for multiple group comparisons, followed by Bonferroni *post hoc* analysis. All *post hoc* effects are presented as q-values corresponding to the corrected statistics from the multiple-comparisons analysis. Linear regression analyses were conducted to assess associations between behavioral metrics and mechanical thresholds. Data are presented as mean ± SEM, with statistical significance defined as p < 0.05. Although sex was included as a factor, it did not reach statistical significance (**Supplemental Figure 1B**); therefore, data from female and male rats were pooled for all analyses.

## Results

### History of Oxycodone IVSA Induces Allodynia Throughout Protracted Abstinence

To evaluate the effects of oxycodone self-administration on mechanical pain sensitivity, rats underwent a standardized intravenous self-administration (IVSA) paradigm^19^ followed by protracted abstinence (**Figure 1**). Mechanical thresholds were assessed at six timepoints post-withdrawal and were compared to age-matched, drug-naïve controls (Naïve). Oxycodone self-administering rats (Oxy IVSA) withstood significantly fewer grams of force during oxycodone withdrawal compared to Naïve, indicative of allodynia, with recovery by 3 weeks (**Figure 2**). There were significant main effects of Oxycodone Intake [F (1, 36) = 9.452; p < 0.01], Time [F (5, 180) = 2.793; p < 0.05], Subject [F (36, 180) = 2.806; p < 0.0001], and Time × Oxycodone Intake [F (5, 180) = 4.129; p < 0.01] (**Figure 2A**). *Post hoc* analysis confirmed significant reductions in force withstood (g) compared to Naïve at 12 hours (q < 0.0001), 1 day (q < 0.001), and 2-weeks (p < 0.01) into oxycodone withdrawal. These results confirmed that oxycodone withdrawal induces protracted mechanical allodynia. No significant difference in mechanical pain threshold was observed between female and male rats throughout protracted abstinence (**Supplemental Figure 1B**).

**Figure 2.**
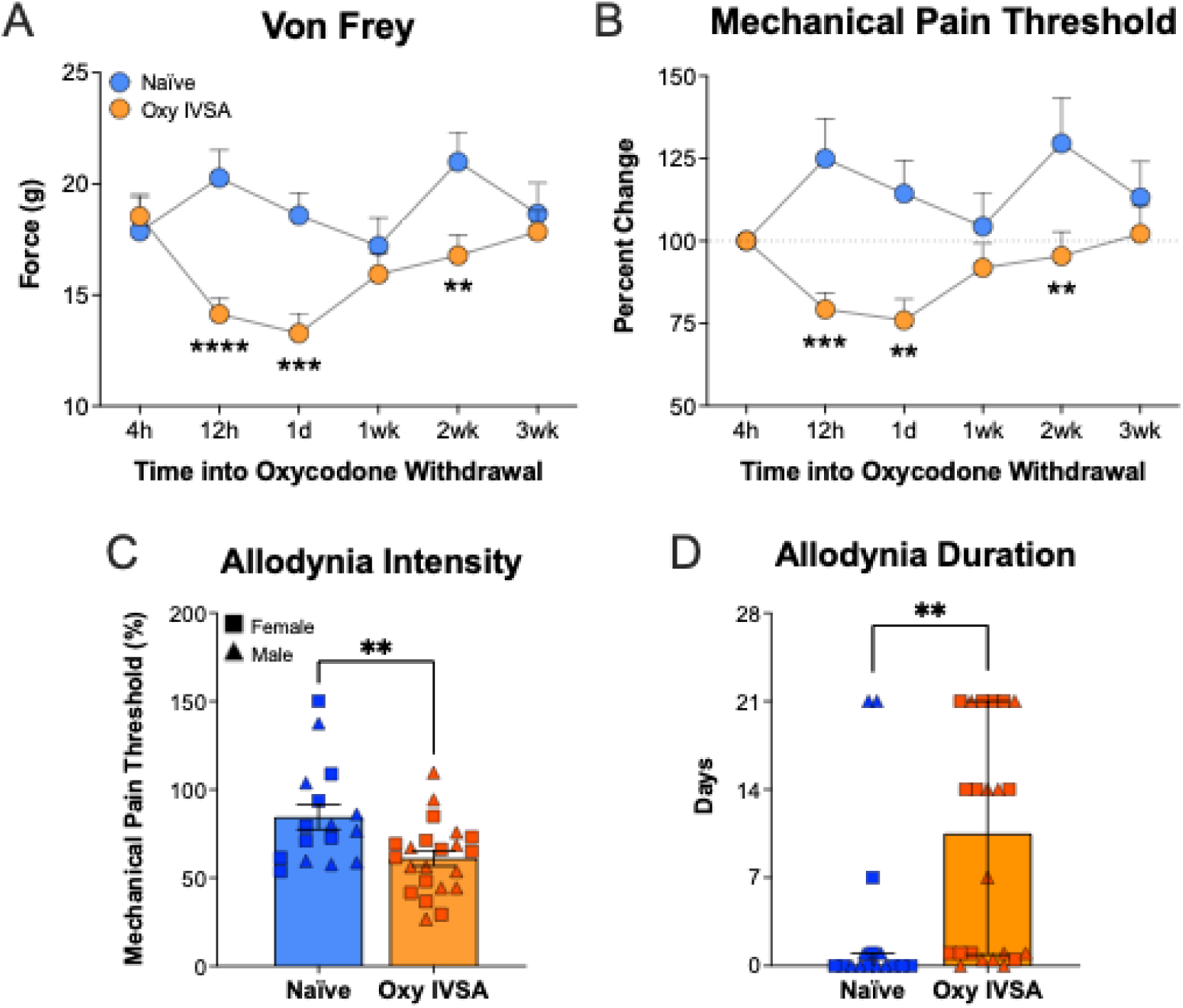
Oxycodone withdrawal-induced allodynia in naïve versus oxycodone self-administering rats. (**A**) Von Frey force withstood (g) and (**B**) mechanical pain threshold (normalized to 4-hours into withdrawal) at six timepoints throughout protracted abstinence. (**C**) Intensity and (**D**) duration of allodynia for each animal, defined as the largest reduction in mechanical pain threshold and the timepoint at which recovery was achieved, respectively. No differences were found between females and males. *p or q < 0.05, **p or q < 0.01, ***p or q < 0.001, ****p or q < 0.0001. n = 22 Oxy IVSA (11/sex) and n = 16 Naïve (8/sex). Oxy IVSA, oxycodone self-administering rats.

To control for within-subject variability in this genetically diverse rat population^57–61^, the mechanical thresholds (in grams) were normalized for each animal and expressed as the percent change from baseline (**Figure 2B**). Baseline was defined as the mechanical threshold at 4 hours into withdrawal. This timepoint was selected for normalization because it was not significantly different from Naïve and, that given the 2 – 4-hour half-life of intravenous oxycodone in rodents^74–76^ (similar to humans^77,78^), residual oxycodone was likely still present in the system, albeit at declining concentrations. This aligns with the onset of early withdrawal signs, which typically emerge 4 – 12 hours after opioid cessation^19,51,79^. Normalization revealed significant main effects of Oxycodone Intake [F (1, 36) = 7.637; p < 0.01], Subject [F (36, 180) = 5.813; p < 0.0001], and Time × Oxycodone Intake [F (5, 180) = 4.387; p < 0.001] (**Figure 2B**). The main effect of Time was p = 0.0566. *Post hoc* analysis again confirmed significant decreases in mechanical pain threshold at 12 hours (q = 0.0001), 1 day (q < 0.01), and 2 weeks (q < 0.01) into withdrawal from oxycodone when compared to Naïve. Allodynia intensity [t(36) = 2.956; p < 0.01] (**Figure 2C**) and duration [Mann-Whitney U = 81, n_1_ = 16, n_2_ = 22, p < 0.01] (**Figure 2D**) were significantly increased and prolonged, respectively, compared to Naïve rats, with on average ∼40% peak reduction in mechanical threshold requiring ∼10 days to recover.

### Stratification Based on Individual Addiction-Like Behaviors

To test if the intensity and duration of allodynia depend on the severity of oxycodone addiction-like behaviors, each animal was categorized based on their individual Addiction Index^19,59,61,80–83^ as High and Low addiction-like subgroups. Four measures were used to generate the Addiction Index for each animal: (1) escalation of oxycodone intake (average number of oxycodone infusions over the last three long access IVSA sessions), (2) motivation to consume oxycodone (LgA PR breakpoint), (3) tolerance to oxycodone’s analgesic effects (difference in tail withdrawal latency, sec; pre- versus post-IVSA), and (4) withdrawal-induced allodynia (mechanical withdrawal threshold, grams of force) (**Figure 3**). High Addiction Index rats had a significantly increased number of oxycodone infusions per session compared to Low Addiction Index rats across short (**Supplemental Figure 2A**) and long access (**Figure 3A**) IVSA. During long access IVSA, High Addiction Index rats showed a consistent and progressive increase in the average number of oxycodone infusions taken across sessions. There were significant main effects of Addiction Index [F (1, 20) = 6.864; p < 0.05], Session [F (13, 260) = 5.114; p < 0.0001], Subject [F (20, 260) = 12.68; p < 0.0001] and Session × Addiction Index [F (13, 260) = 1.922; p < 0.05]. *Post hoc* analysis revealed that on sessions 7 (q < 0.05), 9 (q < 0.01), 12 (q < 0.001), 13 (q < 0.001), and 14 (q < 0.01) High Addiction Index animals self-administered significantly more oxycodone than Low Addiction Index animals (**Figure 3A**). Over the last three sessions specifically (*i.e*., IVSA escalation), High Addiction Index animals had a significantly higher average number of oxycodone infusions than Low Addiction Index animals (p < 0.01; **Figure 3B**).

**Figure 3.**
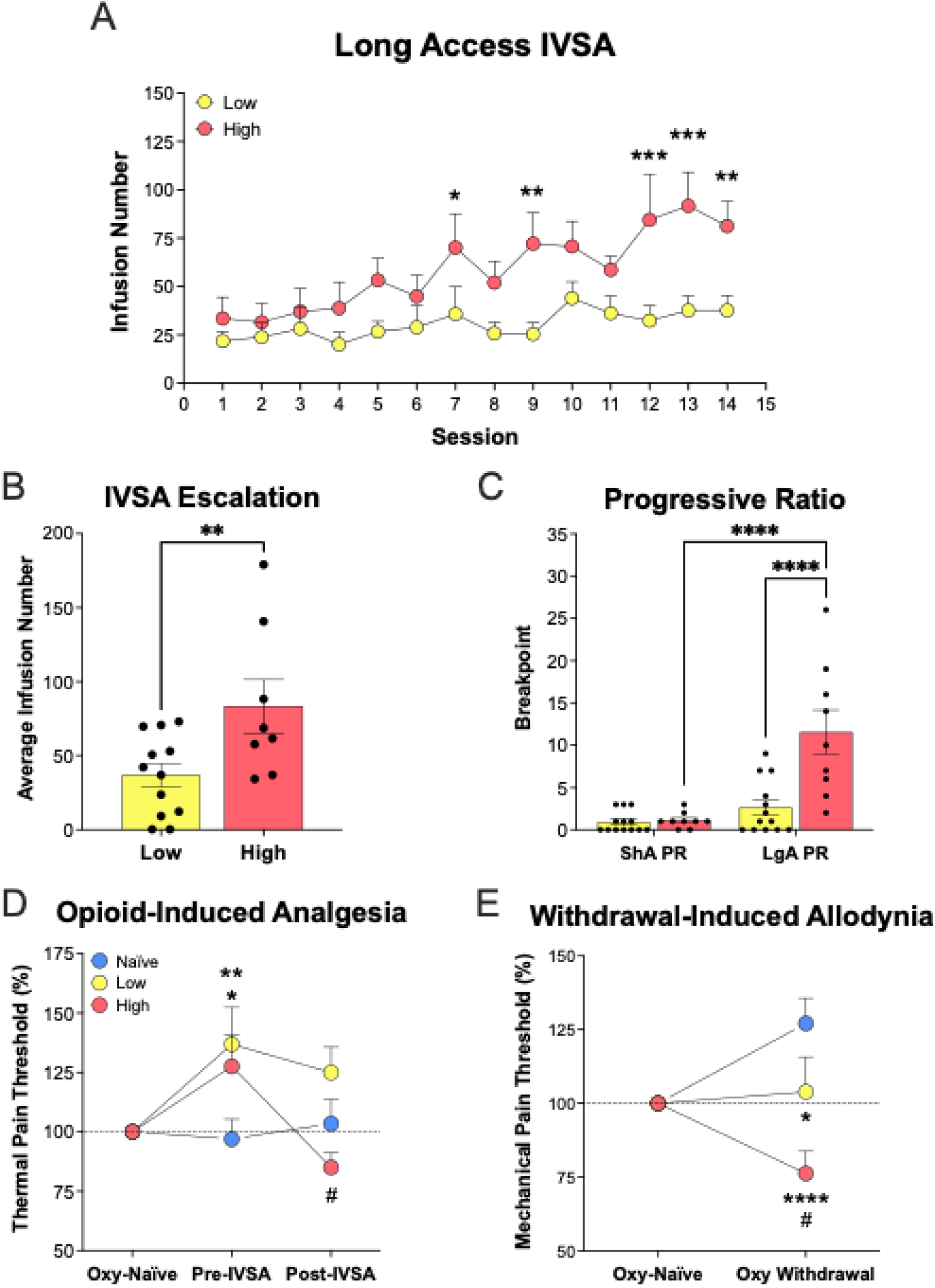
Addiction Index metrics subdivided by index category. (**A**) IVSA infusion number across LgA (12 h/session). (**B**) Oxycodone IVSA esclation (LgA IVSA infusion average over the last three sessions). (**C**) PR responding following ShA and LgA IVSA. (**D**) Opioid-induced analgesia testing conducted before and after LgA IVSA (measure of tolerance and hyperalgesia development), normalized to the oxy-naïve measurement. Oxy IVSA animals were given 450 μg/kg oxycodone i.v. immediately prior to the pre- and post-IVSA analgesia testing. (**E**) Opioid withdrawal-induced allodynia testing conducted 12 h into withdrawal from LgA IVSA and normalized to oxy-naïve measurements. The Naïve group were never exposed to oxycodone and remained as drug-free controls during all behavioral testing. *Post hoc* analysis representations: * significant deviation from Naïve; ^#^significant deviation between High and Low. *^#^p or q < 0.05, **p or q < 0.01, ***p or q < 0.001, ****p or q < 0.0001. n = 9 High (4F/5M), n = 13 Low (7F/6M), and n = 16 Naïve (8/sex). Oxy IVSA, oxycodone-self-administering; Low, Low Addiction Index; High, High Addiction Index; IVSA, intravenous self-administration; ShA, short access; LgA, long access; ShA PR, Progressive Ratio testing following short access IVSA; LgA PR, Progressive Ratio testing following long access IVSA.

PR testing was conducted after both short and long access IVSA (ShA PR and LgA PR, respectively; **Figure 3C**). Analysis showed significant main effects of Addiction Index [F (1, 20) = 13.01; p < 0.001], Long Access IVSA [F (1, 20) = 28.24; p < 0.0001], and Long Access IVSA × Addiction Index [F (1, 20) = 11.71; p < 0.01]. *Post hoc* analysis confirmed significantly higher breakpoints from High Addiction Index rats compared to Low Addiction Index animals during PR responding following long access IVSA (q < 0.0001) but not short access IVSA. High Addiction Index animals further showed a significant increase in breakpoint across PR sessions (q < 0.0001). The results confirm that High Addiction Index rats exhibited more severe addiction-like behaviors across multiple domains.

### Tolerance and Withdrawal Sensitivity Contribute to Addiction Severity

To measure the tolerance to oxycodone’s analgesic effects, tail immersion latency was measured at baseline, pre-, and post-IVSA (**Figure 3D; Supplemental Figure 2B, 2C**). There were significant main effects of Time [F (2, 70) = 4.865; p < 0.05], Subject [F (35, 70) = 1.780; p < 0.05], and Time × Addiction Index [F (4, 70) = 3.429; p < 0.05] (**Figure 3D**). *Post hoc* analysis confirmed oxycodone-induced analgesia at the pre-IVSA timepoint in both High and Low Addiction Index groups (q < 0.05 and q < 0.01, respectively). However, High Addiction Index animals displayed a significant decrease in analgesia at the post-IVSA timepoint compared to Low Addiction Index rats (q < 0.05), who did not show any significant tolerance to the analgesic effects of oxycodone following long access IVSA.

Mechanical pain threshold was measured at two timepoints: oxycodone-naïve and 12 hours into oxycodone withdrawal following long access IVSA (**Figure 3E; Supplemental Figure 2D, 2E**). Naïve animals showed the expected increase in mechanical sensitivity (likely due to animal growth over time, **Figure 1**), however both High and Low Addiction Index animals did not. Low Addiction Index animals showed no change in mechanical pain threshold, and High Addiction Index animals showed a significant decrease from both Naïve and Low Addiction Index groups (**Figure 3E**). There were significant main effects of Addiction Index [F (2, 35) = 6.014; p < 0.01] and Oxycodone IVSA × Addiction Index [F (2, 35) = 6.014; p < 0.01]. *Post hoc* analysis confirmed that both Low and High Addiction Index groups developed allodynia compared to Naïve (q < 0.5 and q < 0.0001, respectively), with High Addiction Index animals having more severe allodynia than Low Addiction Index (q < 0.05), demonstrating the stratification of behavior based on Addiction Index.

### Allodynia Severity and Duration Vary by Addiction-Like Behavior

Reanalysis of the mechanical pain threshold data stratified by Addiction Index was conducted to assess individual differences in allodynia severity. This revealed significant main effects of Addiction Index [F (2, 35) = 4.606; p < 0.05], Time [F (5, 175) = 4.433; p < 0.001], Subject [F (35, 175) = 2.991; p < 0.0001] and Time × Addiction Index [F (10, 175) = 3.302; p < 0.001] (**Figure 4A**). *Post hoc* analysis confirmed that both Low and High Addiction Index animals exhibited significant allodynia at 12 hours (q < 0.0001 and q < 0.01, respectively), 1 day (q < 0.01 and q < 0.001, respectively), and 2 weeks (q < 0.05) post-withdrawal compared to Naïve (**Figure 4A**). However, within-group *post hoc* analysis showed that only High Addiction Index animals exhibited sustained deviations from baseline (12 hours, q < 0.05; 1 day, q < 0.0001; 1 week, q < 0.05; and 3 weeks, q < 0.05). Results remained consistent even after removing the allodynia measure when calculating the Addiction Index (**Supplemental Figure 1A**), demonstrating the robustness of the association between drug taking and drug seeking severity and protracted oxycodone withdrawal-induced allodynia.

**Figure 4.**
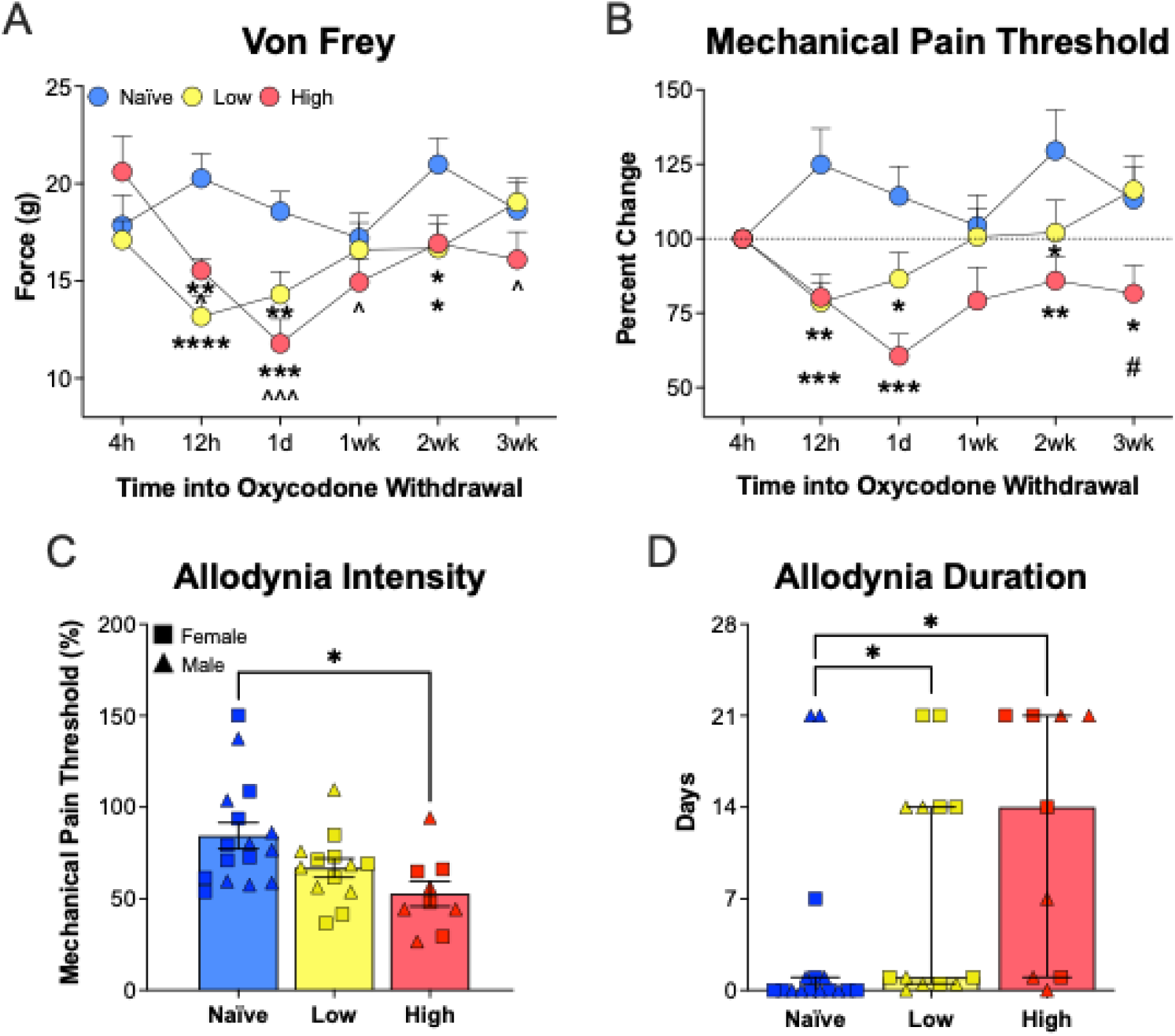
Oxycodone withdrawal-induced allodynia subdivided by Addiction Index. (**A**) Von Frey force (g) and (**B**) mechanical pain threshold (normalized to the 4-hour timepoint) at six timepoints throughout protracted abstinence. (**C**) Intensity and (**D**) duration of allodynia for each animal, defined as the largest reduction in mechanical pain threshold and the timepoint at which recovery was achieved, respectively. *Post hoc* analysis representations: *significant deviation from Naïve; ^#^significant deviation between High and Low; ^significant within-group deviation from baseline timepoint (4 hours). *^#^^p or q < 0.05, **p or q < 0.01, ***^^^p or q < 0.001, ****p or q < 0.0001. n = 9 High (4F/5M), n = 13 Low (7F/6M), n = 16 Naïve (8/sex). Low, Low Addiction Index; High, High Addiction Index.

### Normalization Revealed Divergence in Recovery Timelines

To control for inter-animal variability, mechanical pain thresholds were normalized to the 4-hour post-withdrawal timepoint (**Figure 4B**). Analysis of normalized data revealed significant main effects of Addiction Index [F (2, 35) = 4.984; p < 0.01], Time [F (5, 175) = 2.488; p < 0.05], Subject [F (35, 175) = 5.739; p < 0.0001], and Time × Addiction Index [F (10, 175) = 3.045; p < 0.01]. *Post hoc* comparisons confirmed that High Addiction Index rats showed significant allodynia at 12 hours (q < 0.01), 1 day (q < 0.001), 2 weeks (q < 0.01), and 3 weeks (q < 0.05) post-withdrawal, whereas Low Addiction Index animals had significant allodynia at 12 hours (q < 0.001), 1 day (q < 0.05), and 2 weeks (q < 0.05) only. At 3 weeks, High Addiction Index animals had significantly reduced mechanical threshold compared to both Naïve and Low Addiction Index (q < 0.05). This suggests extended allodynia is experienced in High Addiction Index rats, but not Low Addiction Index rats, persisting for 1 to 3 weeks into abstinence (**Figure 4B**). To control for the effect of potential acute intoxication on the normalization results, the data was alternatively normalized to the drug-naïve baseline measurement (**Figure 1**) and results remained consistent (data not shown). Peak allodynia intensity significantly stratified across groups (p < 0.01; **Figure 4C**) and *post hoc* analysis revealed a significant reduction in High Addiction Index rats relative to Naïve (q < 0.05). Allodynia duration was similarly stratified (p < 0.05; **Figure 4D**), however *post hoc* analysis revealed significantly increased allodynia duration in both groups compared to Naïve (q < 0.05).

### Allodynia Intensity Associates with Prior Oxycodone Intake

Linear regressions were performed to test whether addiction-like metrics predicted mechanical pain sensitivity at each withdrawal timepoint during abstinence (**Figure 5**). Escalation of oxycodone intake (**Figure 5A**), motivation to consume oxycodone (LgA PR; **Figure 5B**), and tolerance to oxycodone’s analgesic effects (**Figure 5C**) were modeled as predictors of the mechanical threshold shifts observed during withdrawal. Long access escalation significantly predicted allodynia severity 1 day into withdrawal across all oxycodone self-administering rats [F (1, 20) = 9.213; p < 0.01] explaining 31.5% of the variance (**Figure 5A**). The LgA PR breakpoint also significantly predicted allodynia severity 1 day into withdrawal [F (1, 20) = 5.545; p < 0.05] explaining 21.7% of the variance (**Figure 5B**). No significant associations were detected between tolerance to oxycodone’s analgesic effects and mechanical thresholds at any withdrawal timepoint (**Figure 5C**). There were also no significant associations detected between early motivation to consume oxycodone (ShA PR; **Supplemental Figure 3A**) or analgesia (**Supplemental Figure 3B**) and mechanical pain thresholds at any withdrawal timepoint.

**Figure 5.**
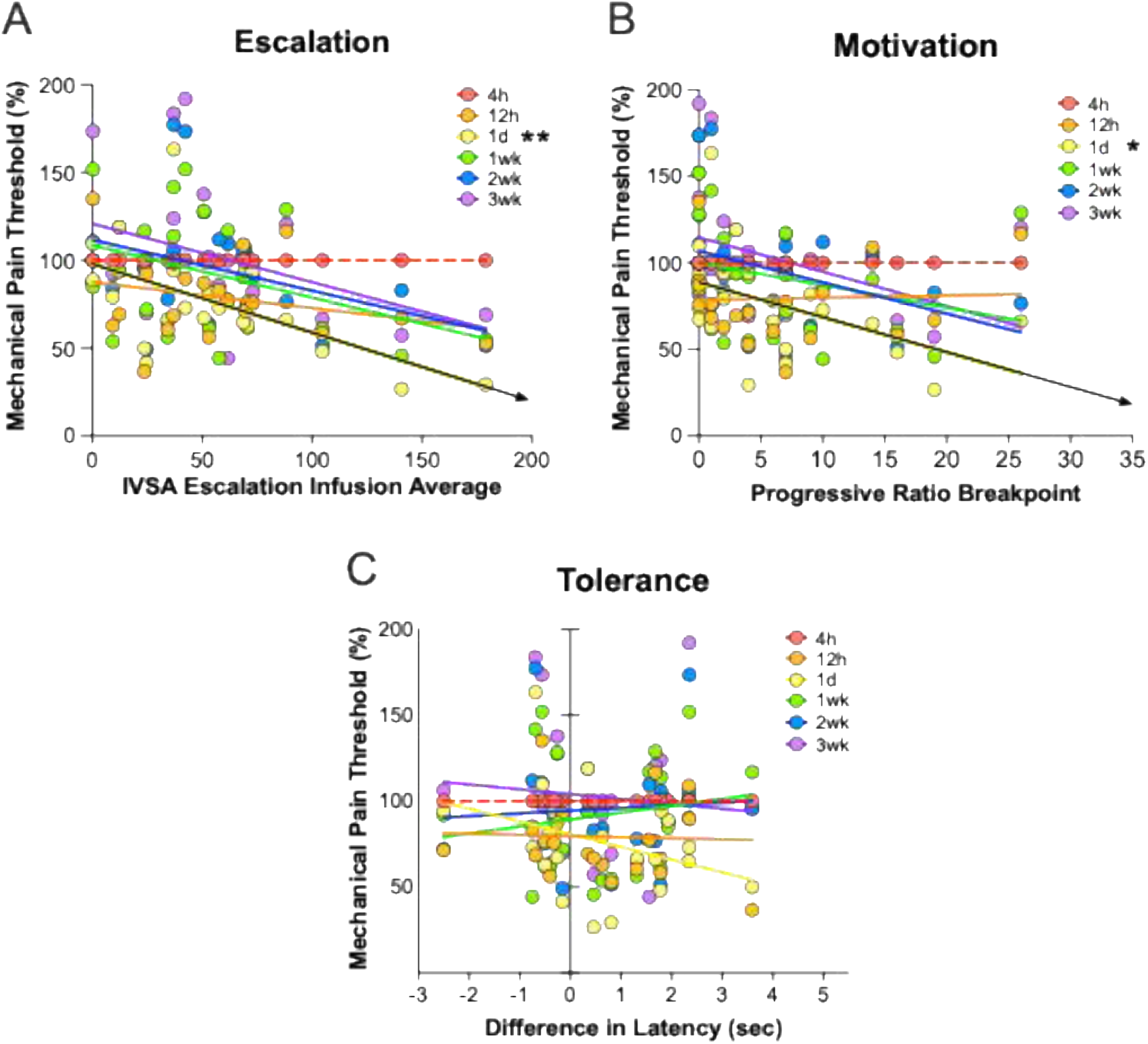
Linear regression of withdrawal-induced mechanical allodynia throughout protracted abstinence versus (**A**) escalation of oxycodone intake (average number of infusions over the last three long access sessions), (**B**) motivation to consume oxycodone (LgA PR breakpoint), and (**C**) tolerance to oxycodone’s analgesic effects (difference in tail-withdrawal latency between pre- and post-IVSA analgesia testing) in all Oxy IVSA rats. Mechanical thresholds at each withdrawal timepoint (4 h, 12 h, 1 d, 1 wk, 2 wk, and 3 wk) were regressed separately for each addiction-like metric. *p < 0.05 and **p < 0.01. n = 22 (11/sex).

## Discussion

Here, we show that a history of oxycodone intravenous self-administration produced prolonged mechanical allodynia throughout protracted abstinence, with significant reductions in mechanical pain threshold persisting from 12 hours to 2 weeks, before recovering by 3 weeks (**Figure 2**). Rats stratified by Addiction Index exhibited clear behavioral divergence, with High Addiction Index animals showing greater escalation of oxycodone intake, stronger motivation to consume oxycodone, increased tolerance to oxycodone’s analgesic effects, and more severe withdrawal-induced allodynia than Low Addiction Index Animals (**Figure 3**). The severity and duration of allodynia differed across addiction-like phenotypes, with High Addiction Index rats displaying both larger and more persistent threshold reductions across abstinence than Low Addiction Index rats (**Figure 4**). Escalation of oxycodone intake and motivation to consume oxycodone predicted allodynia intensity one day into withdrawal, whereas tolerance to oxycodone’s analgesic effects did not, demonstrating that specific addiction-like behaviors – but not analgesic tolerance – predict subsequent pain sensitivity (**Figure 5**). Taken together, these findings identify addiction-like behavioral severity as a strong determinant of withdrawal-induced mechanical allodynia and highlight behavioral predictors that reflect individual vulnerability during opioid abstinence.

### Opioid Withdrawal Induces Prolonged Allodynia in Rats with a History of Oxycodone Self-Administration

The experience of pain is strongly associated with the development and maintenance of Opioid Use Disorder. A history of opioid use lowers the pain perception threshold during withdrawal, with allodynia recovery taking hours to weeks in clinical cases^26,39,42,55,56^. Current data support the stratification of allodynia intensity dependent on the severity of opioid use^19,22,33,34,51^, however, there is no longitudinal characterization of opioid withdrawal-induced allodynia with respect to addiction severity. This report demonstrates that oxycodone IVSA induces prolonged mechanical allodynia during protracted abstinence, with severity and duration dependent on the intensity of the addiction-like behaviors. Significant allodynia was observed in oxycodone self-administering rats at 12 hours, 1 day, and 2 weeks into withdrawal when compared to the Naïve controls, with recovery evident by 3 weeks (**Figure 2A, 2B**). At peak intensity, the pain threshold of Oxy IVSA rats dropped by 40% (**Figure 2C**), taking 10 days to overcome (**Figure 2D**). Of note was the observed variability in pain threshold among Naïve rats. Naïve rats demonstrated normal fluctuations in pain threshold across time (**Figure 2A, 2B**), with the peak reduction in pain threshold (*i.e.,* allodynia intensity) reaching a nadir at 15% below baseline (**Figure 2C**) with no extended duration (**Figure 2D**). However, the range of variability in mechanical pain remained above 100% throughout behavioral testing (**Figure 2B**). Taken together, these findings confirm that oxycodone withdrawal-induced allodynia persists for an extended duration following cessation of use, with 12 hours and 1 day being the most severe timepoints into withdrawal.

### Addiction-Like Behavioral Severity Predicts Allodynia Intensity and Duration

Oxycodone self-administering rats displayed extended allodynia throughout protracted abstinence, however there was a large spread in allodynia severity and duration. To understand the relationship between addiction-like behaviors and allodynia, each rat was subdivided into High and Low based on their individual Addiction Index (**Figure 3**)^19^. High Addiction Index rats self-administered significantly more oxycodone (**Figure 3A, 3B**), displayed greater motivation to consume oxycodone (**Figure 3C**), developed a greater tolerance to oxycodone-induced analgesia (**Figure 3D**), and exhibited more severe oxycodone withdrawal-induced allodynia (**Figure 3E**) when compared to Low Addiction Index rats. This suggests that as oxycodone addiction-like behaviors become more severe over time, oxycodone’s effects paradoxically shift away from analgesic, triggering hyperalgesia and allodynia during withdrawal.

Reassessment of mechanical allodynia throughout protracted abstinence from oxycodone IVSA revealed significant deviations across Addiction Index (**Figure 4**). High and Low Addiction Index rats exhibited similar allodynia at 12 hours into oxycodone withdrawal, however, the High Addiction Index animals then displayed exacerbated allodynia at 1 day, 2 weeks, and 3 weeks into abstinence (**Figure 4B**). Low Addiction Index rats exhibited peak severity at 12 hours, whereas High Addiction Index rats showed a continued decrease in pain threshold through 1 day into oxycodone withdrawal. Further, only High Addiction Index rats displayed significant within-group reductions in mechanical pain threshold from baseline (4 hours; **Figure 4A**). High Addiction Index rats had a pronounced near-50% peak reduction in mechanical threshold during oxycodone withdrawal compared to Naïve (**Figure 4C**), and showed incomplete recovery to baseline by week 3 of withdrawal (**Figure 4D**). In contrast, Low Addiction Index rats experienced about 35% peak reduction in mechanical pain threshold, with recovery to baseline beginning 1 day into abstinence. Thus, addiction-like behavioral severity is directly associated with the intensity and duration of opioid withdrawal-induced allodynia.

### Self-Administration Behavior, But Not Tolerance, Predicts Subsequent Pain Sensitivity

Behavioral markers that predict opioid withdrawal severity have considerable translational value for identifying individuals at elevated risk for severe and prolonged negative withdrawal states. To determine whether addiction-like behaviors were associated with the magnitude of oxycodone withdrawal-induced allodynia, linear regression analyses were performed across several self-administration variables. Escalation of oxycodone intake (average intake during the final IVSA sessions) and motivation to consume oxycodone (LgA PR breakpoint) each predicted the severity of mechanical allodynia at the 1-day withdrawal timepoint (**Figure 5A, 5B**). These findings indicate that higher opioid intake and stronger opioid-directed motivation are associated with more severe early withdrawal pain. The marked divergence in allodynia intensity between High and Low Addiction Index subgroups at this timepoint (**Figure 4B**) further suggests that self-administration behavior may forecast subsequent addiction-like trajectories. As High Addiction Index rats displayed pronounced allodynia for three weeks into protracted abstinence, these markers, if observed in clinical populations, may suggest an extended duration of clinical intervention.

In contrast, no significant linear associations were observed with the tolerance to oxycodone’s analgesic effects (**Figure 5C**), demonstrating that analgesic tolerance does not predict opioid withdrawal-induced pain sensitivity. This dissociation suggests that tolerance arises from distinct neurobiological mechanisms. It may reflect dose-dependent neuroadaptations in modularity circuits^84^ (*e.g.* mesolimbic dopamine system^15,85^, stress circuits^86^, neuroplasticity^87,88^, μ-opioid receptor desensitization^89,90^) or genetic predispositions for differential pain perception^91–94^ and duration^95,96^. Supporting these data, preclinical and clinical evidence show that tolerance reflects receptor-level adaptations during opioid exposure^40–42,97–100^, whereas withdrawal-induced pain emerges from central sensitization and neuroimmune activation during abstinence^101–105^. Individuals may therefore experience severe opioid withdrawal-induced allodynia irrespective of opioid dose or tolerance history^100,106–108^. This is consistent with clinical patterns in which methadone-maintained patients show stable tolerance with minimal withdrawal pain, while others display pronounced allodynia despite lower opioid exposure^28^. These observations support the conclusion that opioid tolerance and withdrawal-induced allodynia are mechanistically and clinically distinct phenomena. Together, these results demonstrate that escalation of opioid use and motivation to consume opioids serve as meaningful behavioral predictors of early withdrawal pain severity and may produce insight into vulnerability for more severe addiction-like outcomes.

### Implications for Clinical Management of Opioid Use Disorder

Opioid withdrawal-induced allodynia may increase relapse vulnerability in individuals with Opioid Use Disorder^42,109–112^. Assessing pain sensitivity and individual phenotypes through Human Genome Wide Association Studies could further support prediction of opioid-risk behavior^113^ and guide personalized treatment strategies based on anticipated pain^94,114^. The prolonged withdrawal-related pain observed in High Addiction Index rats indicates that individuals with more severe addiction may experience extended mechanical sensitivity during abstinence. Although tolerance to oxycodone’s analgesic effects did not predict allodynia in this model, early alterations in analgesia and emerging tolerance remain clinically relevant indicators of heightened vulnerability to protracted abstinence pain and should be evaluated alongside escalation and motivation when assessing individual risk. Identifying early behavioral predictors of addiction vulnerability, such as intake escalation or high motivation to consume opioids, may therefore inform clinical intervention strategies aimed at reducing withdrawal-related discomfort and relapse risk.

### Conclusion

This study demonstrates that opioid withdrawal-induced allodynia varies in both severity and duration according to addiction-like behaviors. High Addiction Index rats exhibited greater and longer-lasting pain sensitivity, which associated with escalation of oxycodone intake and motivation to consume oxycodone. While tolerance to oxycodone’s analgesic effects did not associate with withdrawal-induced allodynia, alterations in analgesic sensitivity still reflected individual differences in addiction-like severity. These findings suggest that mechanical sensitivity during opioid withdrawal may reflect individual vulnerability to addiction-like behavior and that early behavioral metrics may hold predictive value for long-term withdrawal outcomes. Together, these insights may support the development of targeted, personalized interventions for individuals at heightened risk for relapse due to prolonged withdrawal-induced pain.

## Supporting information

Supplemental Results

## Data Availability Statement

The data that support the findings of this study are available from the corresponding author upon reasonable request. Raw and processed data files have been archived on secure institutional serves in compliance with University of California, San Diego data management policies. Access will be granted to qualified researchers for non-commercial use, subject to a data use agreement to ensure responsible handling of sensitive experimental material.

## Declaration of Competing Interests

The authors declare no competing financial interests or personal relationships that could have appeared to influence the work reported in this paper.

## Funding

This work was funded and supported by the National Institute of Health [NIH U01 DA044451, Use of Next-Gen Sequencing to Identify Genetic Variants that Influence Compulsive Oxycodone Intake in Outbred Rats] and the Preclinical Addiction Research Consortium.

## Authors’ contributions

Conceptualization: OG. Methodology: OG, MK, and GdG. Analysis: SP. Investigation: SP, LT, CQ, NT, ES, DO, AM, KB, and SF. Writing: SP, CQ, and NT. Supervision: SP, MK, and OG. Project Administration: SP, ES, MB, LC, MK, GdG, and OG. Funding acquisition: OG.

## Acknowledgements

The authors are grateful to Dr. Solberg Woods and Dr. Palmer for providing the animals used in these experiments. We sincerely thank Dr. Marina Wolf and Dr. Yavin Shaham for providing the initial spart of insight that inspired the foundation of this work. We also thank Angelica Martinez, Selene Bonnet-Zahedi, Ran Qiao, Selen Dirik, Brent Boomhower, Mckenzie Fannon, and all members of the George laboratory for their support.

## Notes

### Competing Interest Statement

The authors have declared no competing interest.

