## Supplemental Results for "Addiction-Like Severity Predicts Prolonged Oxycodone Withdrawal-Induced Allodynia in Genetically Diverse Rats"

*Allodynia is Independent of Sex and Allodynia Outcomes Remain Stable Despite Pain Z-Score Removal.* A second Addiction Index lacking the Pain Z-score was generated and all data reassessed (**Supplemental Figure 1A**). The subdivided results with the Pain Z-score removed were largely comparable to the data including the Pain Z-score, with the mechanical pain threshold throughout withdrawal from oxycodone revealing main effects of Addiction Index [$F(2, 35) = 5.703$; $p < 0.01$], Time [$F(5, 175) = 2.581$; $p < 0.05$], Subject [$F(35, 175) = 5.505$; $p < 0.0001$], and Time × Addiction Index [$F(10, 175) = 2.838$; $p < 0.01$]. *Post hoc* comparisons confirmed that when the Pain Z-score was omitted, High Addiction Index animals showed significant allodynia at 12 hours ($q < 0.001$), 1 day ($q < 0.001$), and 2 weeks ($q < 0.01$) post-withdrawal, with trending significance at 3 weeks post-withdrawal ($p = 0.0605$) when compared to Naïve. Low Addiction Index animals (lacking the Pain Z-score) showed significant allodynia at 12 hours ($q < 0.01$), and trending significance for allodynia at 1 day ($q = 0.0557$) when compared to Naïve. Further, there was trending *post hoc* significance between High and Low Addiction Index animals at 1 day ($q = 0.0557$) and 3-weeks ($q = 0.0605$) post-withdrawal (when pain was excluded from the Addiction Index). As the data omitting the Pain Z-score was largely comparable to its inclusion, the Pain Z-score was included in all analyses as it strengthens the Addiction Index.

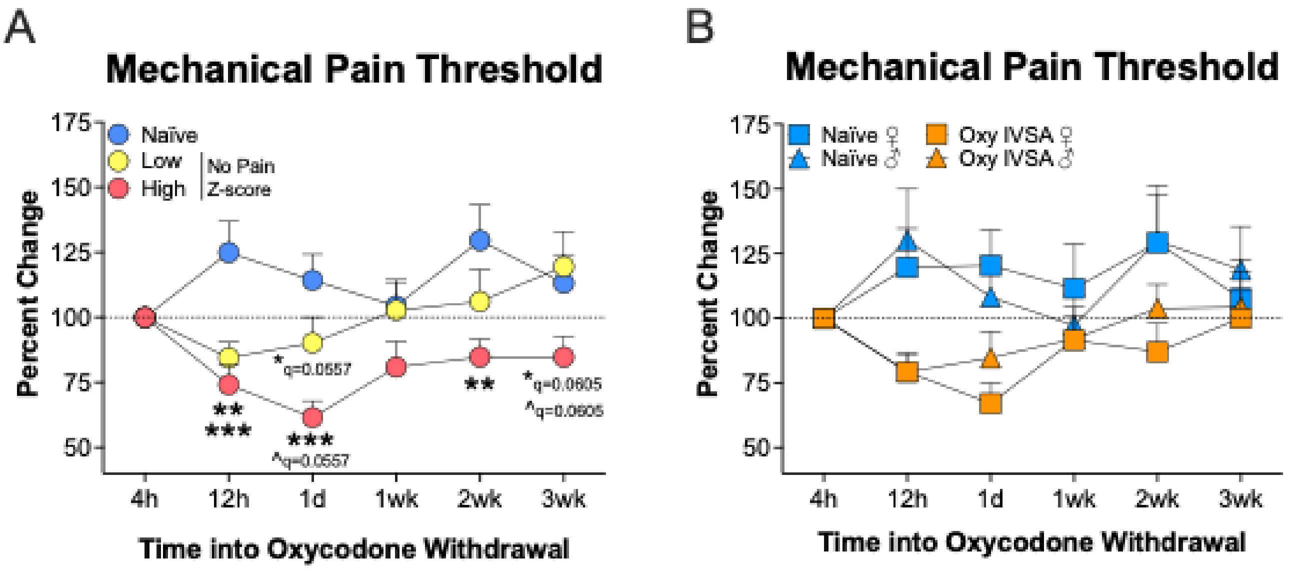


**Supplemental Figure 1**. Behavioral phenotypes are not influenced by exclusion of Pain Z-score from the Addiction Index or sex. (**A**) Mechanical pain threshold (normalized to the 4-hour timepoint) at six timepoints throughout protracted abstinence, subdivided by an alternate Addiction Index that lacked the Pain Z-score. (**B**) Mechanical pain threshold throughout withdrawal comparing Naïve and Oxy IVSA rats, broken down by sex. *Post hoc* analysis representations: *significant deviation from Naïve; ^significant within-group deviation from baseline timepoint (4 h). *p or q < 0.05, **p or q < 0.01, and ***p or q < 0.001 compared to naïve. n = 22 Oxy IVSA (11/sex), n = 11 High (6F/5M), n = 11 Low (6F/5M); n = 16 Naïve (8/sex).

There was no significant effect of sex identified across the mechanical pain testing timeline (**Supplemental Figure 1B**). Therefore, female and male animals were pooled for all analyses. Animals were subdivided into High and Low Addiction Index based on their respective Addiction Index Z-scores and their behavior compared. It should be noted that the Pain Z-score used in the overall Addiction Index is measured via Von Frey. It was therefore necessary to ensure the Pain Z-score did not significantly bias the subsequent results.

*Rats Stratify Based on Individual Addiction-Like Behaviors.*

During the short access IVSA training sessions, High Addiction Index rats consistently had a significantly higher number of oxycodone infusions when compared to Low Addiction Index rats (**Supplemental Figure 2A**). There were significant main effects of Addiction Index [$F(1, 20) = 5.252$; $p < 0.05$] and Subject [$F(20, 60) = 3.035$; $p < 0.001$]. Tail immersion responses following a history of oxycodone IVSA were compared to Naïve (**Supplemental Figure 2B**). Three measurements were used: baseline (no oxycodone), analgesia (450 µg/kg oxycodone with no history of oxycodone IVSA), and tolerance (450 µg/kg oxycodone following a history of oxycodone IVSA; **Figure 1**). It should be noted that Naïve rats were tested at all three timepoints but were never exposed to oxycodone (*i.e.,* not 'analgesia' or 'tolerance' measures for control). There were significant main effects of Oxycodone Intake [$F(1, 36) = 6.642$; $p < 0.05$] Subject [$F(36, 72) = 2.822$; $p < 0.0001$], and Time x Oxycodone Intake [$F(2, 72) = 5.486$; $p < 0.01$]. *Post hoc* analysis revealed a significant increase in analgesia following oxycodone administration with no previous history of oxycodone exposure (pre-IVSA; $q = 0.0001$) (**Supplemental Figure 2B**). The animals were subdivided by Addiction Index (**Supplemental Figure 2C**). There were significant main effects of Time [$F(2, 70) = 3.687$; $p < 0.05$], Subject [$F(35, 70) = 3.117$; $p < 0.0001$], and Time x Addiction Index [$F(4, 70) = 4.770$; $p < 0.01$]. Additionally, there was a near-significant main effect of Addiction Index ($p = 0.0516$). *Post hoc* analysis revealed significant analgesia in both High and Low Addiction Index groups when compared to Naïve control ($q < 0.001$) (**Supplemental Figure 2C**). When considering within-group *post hoc* analysis, while Low Addiction Index animals showed significant variation between baseline and both analgesia ($q < 0.01$) and tolerance ($q < 0.05$), High Addiction Index rats only showed variability between analgesia and tolerance ($q < 0.01$) (**Supplemental Figure 2C**). There was, however, a near-significant increase in analgesia between baseline and analgesia ($q = 0.0697$).

Mechanical sensitivity data used to generate the Pain Z-score was calculated from two timepoints (**Figure 1**): oxycodone-naïve and 12 hours into oxycodone withdrawal (following long-access IVSA)

(**Supplemental Figure 2D**). There was a significant main effect of Subject [$F(36, 36) = 1.969$; $p < 0.05$] and Time x Oxycodone intake [$F(1, 36) = 8.293$; $p < 0.01$]. There was a trending main effect of Oxycodone Intake ($p = 0.0538$). *Post hoc* analysis revealed a significant decrease in the grams of force withstood by the Oxy IVSA group during withdrawal when compared to Naïve ($q < 0.01$) (**Supplemental Figure 2D**). The animals were then subdivided by Addiction Index (**Supplemental Figure 2E**). There was a significant main effect of Subject [$F(35, 35) = 2.144$; $p < 0.05$] and Time x Addiction Index [$F(2, 35) = 6.010$; $p < 0.01$]. *Post hoc* analysis revealed significantly lower levels of force withstood by both High and Low Addiction Index rats when compared to Naïve ($q < 0.01$) (**Supplemental Figure 2E**).

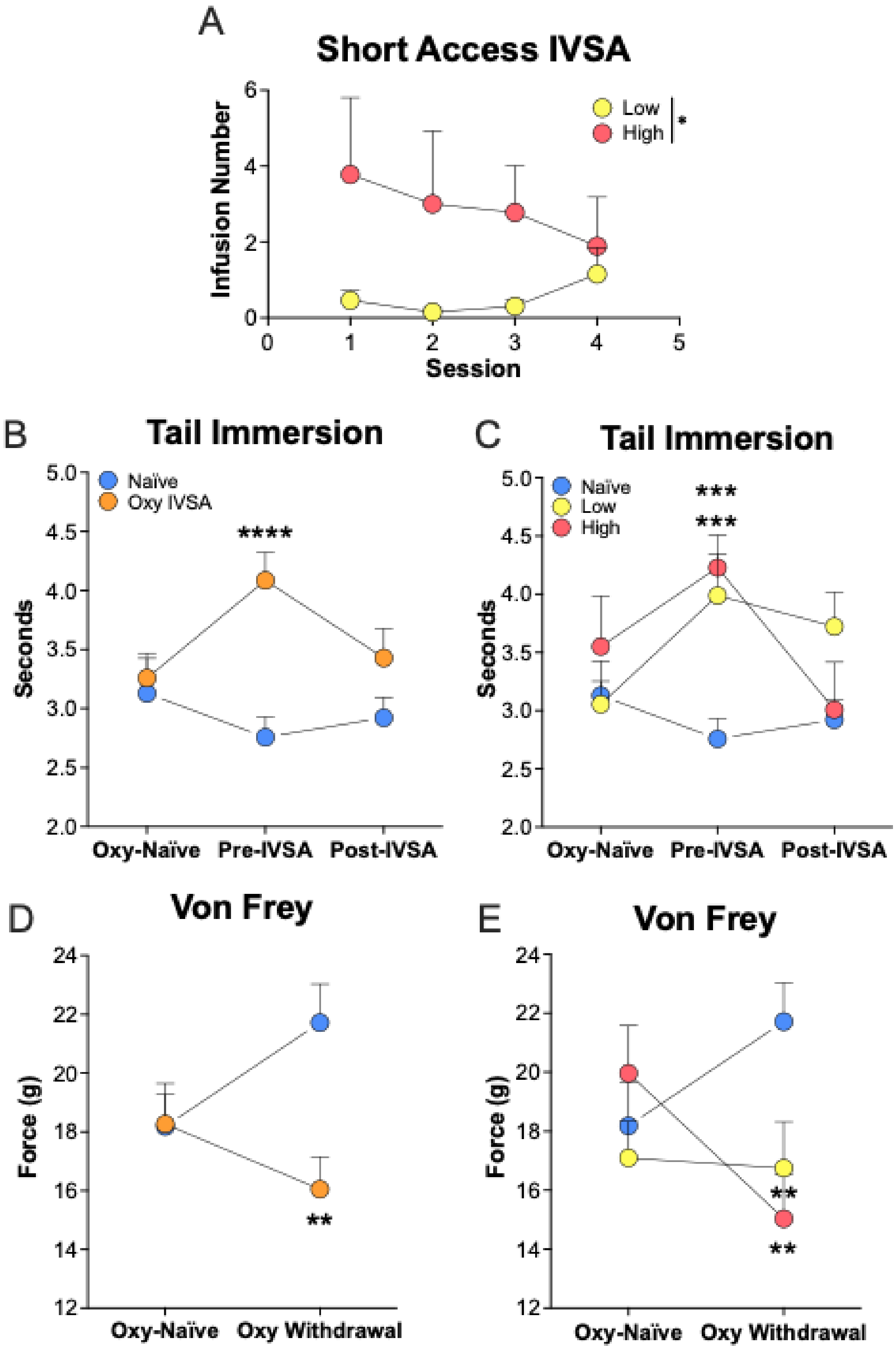


**Supplemental Figure 2**. Supporting behavioral data for Addiction Index generation. (**A**) Short access IVSA behavior subdivided by Addiction Index (2 h/session). Tail immersion time comparing (**B**) Naïve and Oxy IVSA rats, and (**C**) Naïve, Low, and High Addiction Index rats across three timepoints during IVSA behavioral testing. Oxy IVSA animals were given 450 µg/kg oxycodone i.v. immediately prior to pre- and post-IVSA testing. Von Frey testing comparing (**D**) Naïve and Oxy IVSA rats, and (**E**) Naïve, Low, and High Addiction Index rats at two timepoints (oxycodone-naïve and 12 h into withdrawal following LgA

IVSA). Naïve rats were never exposed to oxycodone and remained as drug-free controls during all behavioral testing. *p or q < 0.05, **p or q < 0.01, ***p or q < 0.001, ****p or q < 0.0001. n = 22 Oxy IVSA (11/sex); n = 9 High (4F/5M), n = 13 Low (7F/6M); n = 16 Naïve (8/sex). Oxy IVSA, oxycodone self-administering rats.

*Allodynia does not Associate with Early Motivation or Analgesia.*

Linear regression analysis revealed no significant associations between mechanical pain threshold and early motivation (ShA PR; **Supplemental Figure 3A**) or analgesia (latency for tail withdrawal pre-IVSA; **Supplemental Figure 3B**) at any withdrawal timepoint.

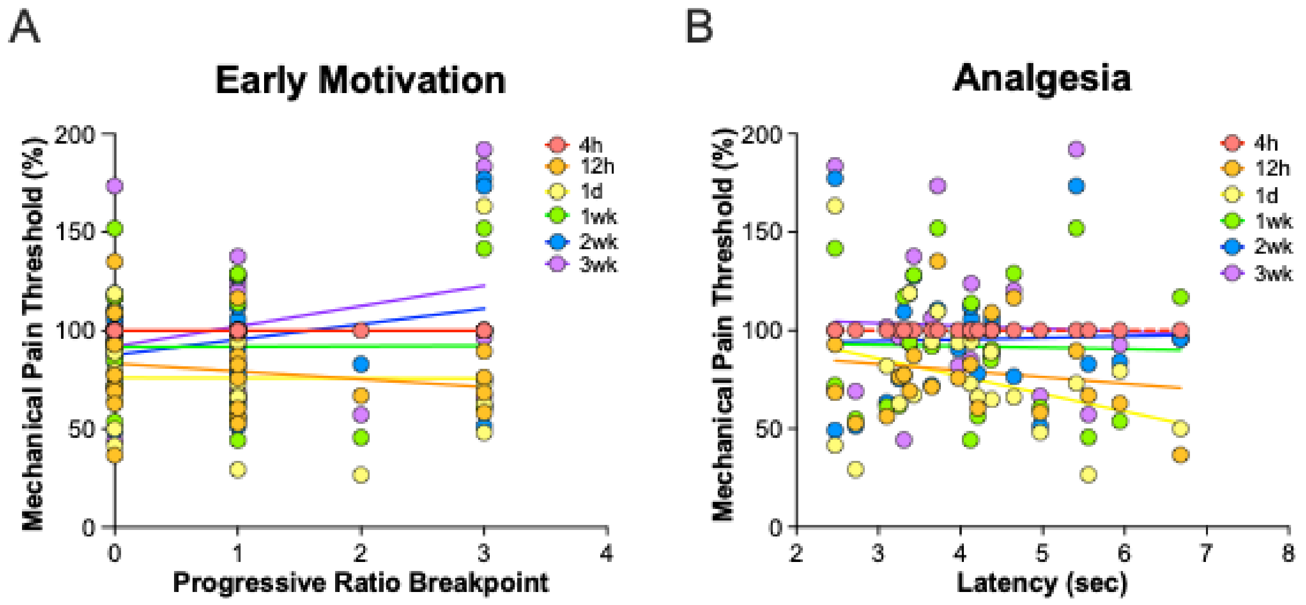


**Supplemental Figure 3**. Protracted allodynia is not associated with early motivation or analgesic tolerance. (**A**) Linear regression of mechanical pain threshold throughout abstinence versus early motivation (ShA PR breakpoint). (**B**) Linear regression of mechanical pain threshold versus opioid-induced analgesia. n = 22 (11/sex).
